# Seeing versus Knowing: The Temporal Dynamics of Real and Implied Colour Processing in the Human Brain

**DOI:** 10.1101/369926

**Authors:** Lina Teichmann, Tijl Grootswagers, Thomas Carlson, Anina N. Rich

**Affiliations:** Perception in Action Research Centre & Department of Cognitive Science, Macquarie University, Sydney, Australia; ARC Centre of Excellence in Cognition & its Disorders, Macquarie University, Sydney, Australia; School of Psychology, University of Sydney, Australia; Centre for Elite Performance, Training and Expertise, Macquarie University, Sydney, Australia

## Abstract

Colour is a defining feature of many objects, playing a crucial role in our ability to rapidly recognise things in the world around us and make categorical distinctions. For example, colour is a useful cue when distinguishing lemons from limes or blackberries from raspberries. That means our representation of many objects includes key colour-related information. The question addressed here is whether the neural representation activated by *knowing* that something is red is the same as that activated when we *actually see* something red, particularly in regard to timing. We addressed this question using neural timeseries (magnetoencephalography, MEG) data to contrast real colour perception and implied object colour activation. We applied multivariate pattern analysis (MVPA) to analyse the brain activation *patterns* evoked by colour accessed via real colour perception and implied colour activation. Applying MVPA to MEG data allows us here to focus on the temporal dynamics of these processes. Male and female human participants (N=18) viewed isoluminant red and green shapes and grey-scale, luminance-matched pictures of fruits and vegetables that are red (e.g., tomato) or green (e.g., kiwifruit) in nature. We show that the brain activation pattern evoked by real colour perception is similar to implied colour activation, but that this pattern is instantiated at a later time. These results suggest that a common colour representation can be triggered by activating object representations from memory and perceiving colours.

## Introduction

Throughout our lives, we learn statistical regularities about objects in our environment. We acquire knowledge about their typical perceptual features, which motor actions are required to interact with them, and in which context they usually appear. For example, we know that a tomato is round and red, we can eat it and it appears in the wider context of food. Our neural representations of objects therefore need to encompass a conceptual combination of these learnt attributes spanning from perception to action and semantic knowledge (A. Martin, Haxby, Lalonde, Wiggs, & Ungerleider, 1995). The activation of object representations is likely to involve a widespread, distributed activation of several brain regions (Patterson, Nestor, & Rogers, 2007) with some brain areas responding preferentially to object colour (e.g., Seymour, Williams, & Rich, 2015). Several neuroimaging studies have compared perceiving colour and accessing object-colour knowledge from memory, finding evidence that similar brain areas are involved in these two processes (e.g., Bannert & Bartels, 2013; A. Martin et al., 1995; Vandenbroucke, Fahrenfort, Meuwese, Scholte, & Lamme, 2014). Using magnetoencephalography (MEG), we look at the neural timecourse of ‘real’ (by which we mean ‘induced by wavelengths of light’) colour perception versus implied object-colour activation from memory.

Associations between objects and typical or *implied* colours are acquired through experience (Bartleson, 1960; Hering, 1920) and are activated effortlessly and involuntarily (Bramão, Faísca, Petersson, & Reis, 2010; Chiou & Rich, 2014). The activation of object-colour knowledge is part of the dynamic interaction between perceptual processes and activation of prior conceptual knowledge to evaluate sensory input (Collins & Olson, 2014; Engel, Fries, & Singer, 2001; Goldstone, de Leeuw, & Landy, 2015). One of the central questions is how object-colour knowledge interacts or overlaps with colour representations generated by external stimuli. There is behavioural evidence that object-colour knowledge can influence colour perception. Hansen, Olkkonen, Walter, & Gegenfurtner (2006) found that participants overcompensated for implied colours when they were asked to change the colour of colour-diagnostic objects to be achromatic. For example, a banana would be adjusted towards the blue side of grey, showing the influence of the implied colour yellow. Similarly, Witzel (2016) showed that participants selected an image of an object as achromatic more often when its colour was modified to be the opposite of its implied colour (e.g., a bluish-grey banana). These results suggest that colour perception can be influenced by previously learnt object-colour associations (see Firestone and Scholl, (2016) for debates about the extent to which activation of colour from memory is identical to colour perception). Brain-imaging data, recorded with functional magnetic resonance imaging (fMRI), suggest that brain activation corresponding to implied object colour activation shares characteristics with real colour perception: Retrieving the knowledge that a banana is yellow activates brain areas in or around the V4 complex, which is involved in colour perception (Bannert & Bartels, 2013; Barsalou, Simmons, Barbey, & Wilson, 2003; Chao & Martin, 1999; Rich et al., 2006; Simmons et al., 2007; Vandenbroucke et al., 2014). This suggests that activation of implied colour rests on a similar neural architecture as real colour perception.

These results suggest that similar brain areas are active when perceiving colour and accessing implied colour, which may drive the behavioural interactions between the two (e.g., Hansen et al., 2006). Real colour activations occur very early in visual processing, whereas implied colour presumably is only activated once the object is processed at a higher level. Hence, there could be a temporal delay for activity driven by implied colour in comparison to activity driven by perceived colour. As the signal measured by fMRI is slow, it is not a suitable method to distinguish fine temporal differences between real and implied object colour processing. In the current study, we use multivariate pattern analysis (MVPA) on MEG data (Grootswagers, Wardle, & Carlson, 2017) to compare the brain activation patterns evoked by colour perception and implied object colour activation. MEG has fine temporal resolution, and with MVPA we can detect patterns across the sensors at each time point that are reliable enough to train an algorithm to classify different categories of stimulus. Here, we use these methods to test whether a classifier trained on ‘real colour’ can successfully decode ‘implied colour’. Such cross-generalisation can only occur if there is sufficient similarity in the neural signals. This approach enables us to contrast the temporal dynamics of real and implied colour processing, shedding light on the interaction between perceptual processing and activation of object representations.

## Methods

### Participants

20 healthy volunteers (12 female, mean age = 27.6 years, SD = 6.6 years) completed the study. All participants reported normal or corrected-to-normal vision including normal colour vision. Participants gave informed consent before the experiment and were reimbursed with $20/hour. During the MEG recording, participants were asked to complete a target-detection task to ensure they were attentive. Two participants performed more than three standard deviations below the group mean on this task, suggesting they did not pay attention to the stimuli, and were therefore excluded from analysis, leaving 18 participants in total. The study was approved by the Macquarie University Human Research Ethics Committee.

### Procedure

While lying in the magnetically shielded room (MSR) for MEG recordings, participants first completed a colour flicker task (Kaiser, 1991) to equate the coloured stimuli in perceptual luminance. Then they completed the main target-detection task. We used only two colours to increase the power of our analysis. If there are luminance differences between the colour categories, the classifier can use this strong signal to discriminate the categories instead of relying on colour. While previous studies have shown the greatest behavioural effects for colours along the daylight axis (yellow-blue, Hansen et al., (2006)), these are not feasible colours for the current design: equiluminant blue and yellow stimuli no longer look clearly blue and yellow. We chose red and green as the two colours as they can be matched for luminance, and we included varying exemplars of these two hue categories to ensure any potential remaining slight differences in luminance could not be used by a classifier to distinguish the colour categories.

### Colour Flicker Task

In the colour flicker task, participants were presented with red and green circles (5×5 degrees visual angle) in the centre of the screen. The colours alternated at a rate of 30Hz. Participants completed 2 runs of 5 trials each. In each trial, one red-green combination was used. The red colour was kept consistent throughout each trial while participants were asked to use a button box to adjust the luminance level of green and report when they perceived the least amount of flickering. The HSV (hue, saturation, value) values for each green shade were then recorded. This procedure was repeated in the second run. The average HSV values between the two runs was then computed, yielding five shades of red and green equated for perceptual luminance. Using different shades of red and green which were each equated for perceptual luminance minimises the degree that any luminance difference between the categories could influence the results (see Table 1 [supplementary materials] summarising individual HSV values used).

### Target-Detection Task

In the main target-detection task (Figure 1A), participants completed eight blocks of 440 trials each. There were two different types of blocks: *implied colour* and *real colour*. Block types alternated for each participant and the overall order was counterbalanced across participants. In the *implied colour* blocks, participants viewed luminance-equated (SHINE toolbox Willenbockel et al. (2010)) grey-scale images of colour diagnostic objects (see Figure 1A). Equating the overall luminance ensures that differences in MEG response patterns are not caused by luminance differences between the ‘usually red’ and ‘usually green’ categories. To increase variability in the stimulus set, half the stimuli depicted a single item on the screen (e.g., one strawberry) and the other half were multiple, partially overlapping items (e.g., three strawberries). Having several different stimuli in each category helps to minimise the influence of low-level features such as edges and shapes on the results. In the *real colour* blocks, participants viewed five different abstract shapes. Each shape was filled in one of the red and green shades which had been equated for perceptual luminance with the colour flicker task. Each shape occurred equally often in red and green. To match the stimuli presented in the *implied colour* block, half of the shapes were single shapes (e.g., one square) on the screen while the other half consisted of partially overlapping shapes (e.g., three squares). All stimuli (objects and shapes) contained the same number of pixels (Figure 1A).

**Figure 1:**
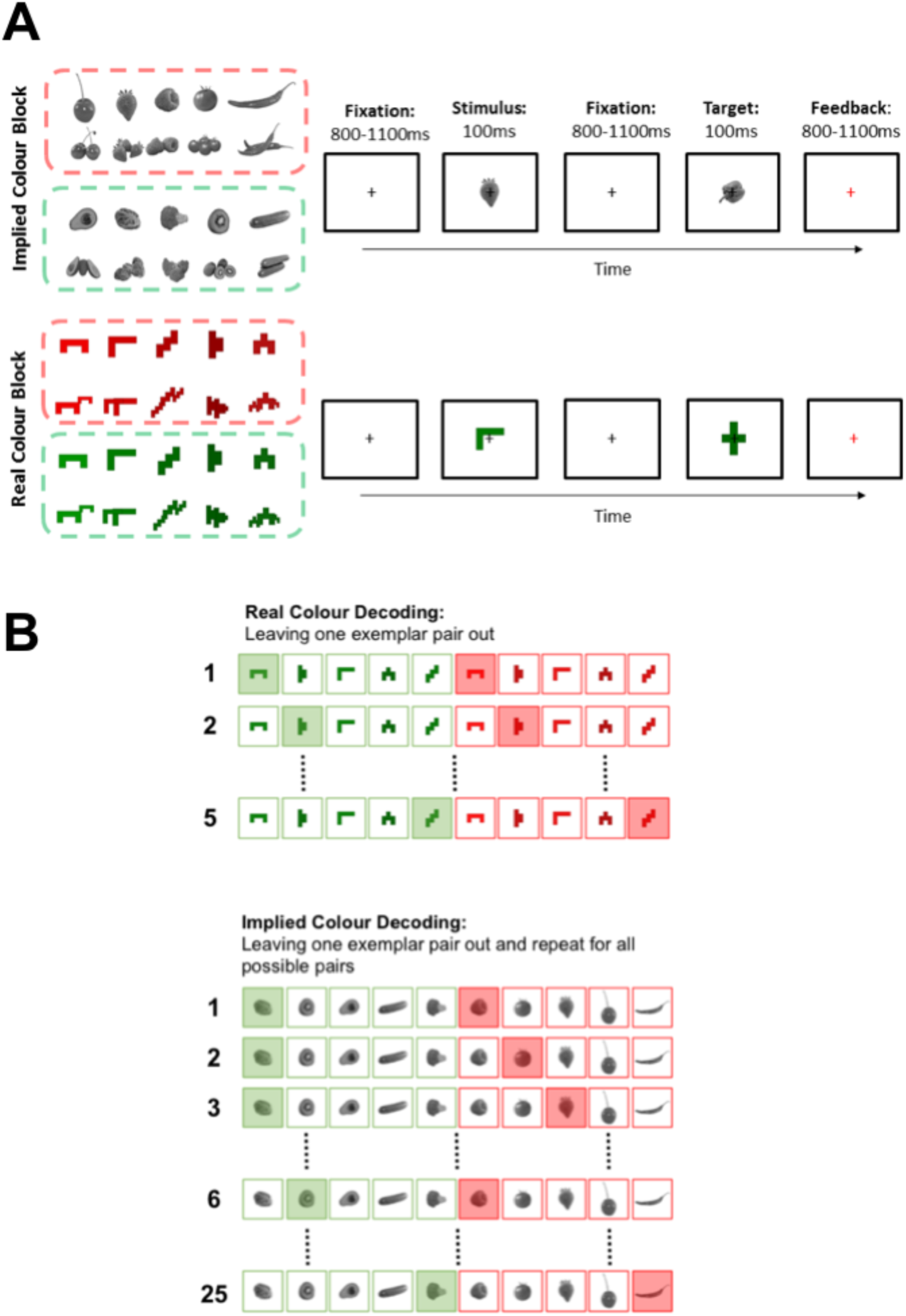
(A) Target-detection task and stimuli for both implied colour (top panel) and real colour (bottom panel) blocks. Participants were asked to press a button as soon as they saw the target (capsicum or cross). If they pressed the button after a target (correct detection) the fixation cross would turn green briefly, if they missed the target it would turn red. (B) Cross-validation approach used for real (top) and implied (bottom) colour decoding analyses. Every row shows which trials were used for the training set (clear) and which trials were used for the testing set (shaded). Trials with the same exemplar are never in the training and the testing set. In the real colour decoding analysis, we split the data in 5 different ways, always leaving one pair of the same shape with matched luminance out. In the implied colour decoding analysis, we split the data in 25 different ways, leaving all possible exemplar pairs out once. The classification accuracy is an average of the classifier performance for each of these divisions.

In both block types, presentation location varied randomly by ∼1 degree visual angle around the central fixation cross. Changing the spatial location of the stimulus images adds variability to the retinal image, again reducing the influence low-level visual features have on the results. Participants were asked to press a button when they saw an image of the target shape (cross) or target object (capsicum). Every block had 40 target trials. All target trials were discarded before the MEG data analysis. On average, participants detected 98.51% (SD = 0.013%) of target stimuli.

### Apparatus and Pre-processing

The stimuli had a visual angle of approximately 5 × 5 degrees and were projected onto a translucent screen mounted in the magnetically shielded room. The stimulus display system used was an Epsom projector (EB-G7400U) with a refresh rate of 60Hz. The projector display size had approximately 10.02 ×19.9 degrees visual angle and the resolution of the display was set to 1280 ×720 pixels. Stimuli were presented using MATLAB with Psychtoolbox extension (Kleiner et al., 2007; Pelli, 1997). The neuromagnetic recordings were obtained with a whole-head axial gradiometer MEG (KIT, Kanazawa, Japan), containing 160 axial gradiometers. The frequency of recording was 1000Hz. FieldTrip (Oostenveld, Fries, Maris, & Schoffelen, 2011) was used to pre-process the data. We used a low-pass filter of 200Hz and a high-pass filter of 0.03Hz online. Stimulus onsets were determined with a photodiode that detected light change when a stimulus came on the screen. Trials were epoched from –100 to 800ms relative to stimulus onset and downsampled to 200Hz (5ms resolution). All target trials were removed. We performed no further preprocessing steps (e.g., channel selection, artefact correction), leaving our data in the rawest possible form. This choice was motivated by recent work showing that traditional preprocessing choices can introduce artefacts in the data that have a strong effect on multivariate analyses (van Driel, Olivers, & Fahrenfort, 2019).

### Decoding Analysis

We conducted four separate decoding analyses using linear discriminant classifiers (LDA) implemented in CoSMoMVPA (Oosterhof, Connolly, & Haxby, 2016). First, to test whether we can decode perception of red versus green, we analysed the data from the real colour (shape) blocks. We tested whether we could decode the colour of our abstract shapes for each person. The classifier was trained on distinguishing the activity patterns evoked by red versus green shapes at each timepoint using 80% of the real colour data. We then used the classifier to predict the colour of each stimulus at every timepoint in the remaining 20% of the real colour data. To divide the data into training and testing set, we used an independent exemplar cross-validation approach (Carlson, Tovar, Alink, & Kriegeskorte, 2013), leaving out one exemplar pair with matched luminance (e.g., red and green L-shape, matched for perceptual luminance). This process was repeated over all folds so that each exemplar pair was in the training and the testing set once (5-fold cross-validation). Hence, the colours in each fold were balanced (Figure 1B).

Second, to assess whether we can decode implied colour from grey-scale objects, we trained a classifier to distinguish trials of grey-scale objects that are associated with red versus green. As in the analysis described above, we used an independent cross-validation approach and trained the classifier on 80% of the implied colour data and tested its performance on the remaining 20% of implied colour data. Because the greyscale objects in the red and green condition varied in more ways than just their implied colours, we left out both possible exemplar pairs for each object in the implied colour decoding analysis to minimise the degree to which visual features such as shape would be used by the classifier. We selected trials based on label for both colour categories (e.g., all strawberry and kiwifruit trials). Note that there were two instances of each stimulus (e.g., an image of one strawberry and an image of three strawberries) and these were considered the same object for the leave-one-out procedure. We trained our classifier to distinguish between activity patterns evoked by all stimuli except the selected stimuli and tested its performance on the left-out trials. We repeated this process to have every possible combination of green and red objects used once as the testing set (25-fold cross-validation), and report the average classification performance over all these combinations (Figure 1B). Although the independent cross-validation approach reduces the risk of features other than implied colour driving the effect, we still have to be cautious with the interpretation as there may be overall low-level differences across all the red and green objects. This is unavoidable when using natural objects.

Last, we conducted a cross-decoding analysis across the two different block types, training the classifier on all real colour trials and testing on all implied colour trials. This cross-decoding analysis is highly conservative as *everything* about the stimuli differs between real colour and object colour trials, the only potential link is the implied colour of the objects to the real colour of the abstract shapes. If there are any low-level differences in the real colour decoding other than chromaticity (e.g., overall luminance difference), this would only decrease the likelihood of finding significant cross-generalisation to the implied colour trials. In addition, any differences in between the greyscale objects cannot drive an effect in the cross-decoding analysis, as the classifier is trained to distinguish the real colour shapes which are the same in the red and the green condition.

It is possible that a similar pattern is elicited by the two colour types but it occurs at different times, thus, we may not see it in a direct cross-decoding analysis. We therefore also conducted a time-generalisation analysis (Carlson, Hogendoorn, Kanai, Mesik, & Turret, 2011; King & Dehaene, 2014), training the classifier at each timepoint on the real colour trials and then testing on each timepoint in implied colour trials, yielding an accuracy score for all pairs of training and testing timepoints. This technique allows us to test for similar activation patterns that do not occur at the same time.

### Statistical Tests Classification Analyses

To assess whether the classifier could distinguish between red and green trials significantly above chance, we used random effects Monte-Carlo cluster statistic (Maris & Oostenveld, 2007) using Threshold Free Cluster Enhancement (TFCE, Smith & Nichols, 2009) as implemented in CoSMoMVPA (Oosterhof et al., 2016). The TFCE statistic represents the support from neighbouring time points, allowing optimal detection of sharp peaks, as well as sustained weaker effects. First, a permutation test was conducted by swapping the labels of complete trials and then we re-ran the analysis on the data with the shuffled labels. This was repeated 100 times per participant to generate subject-level null-distributions. Second, Monte-Carlo sampling was used to create a group-level null-distribution consisting of 10,000 shuffled label permutations for the time-resolved decoding, and 1000 for the time-generalisation analysis (to limit computation time). Third, these group-level null-distributions were transformed into TFCE statistics (Smith & Nichols, 2009). To correct for multiple comparisons, we then selected the maximum TFCE value across time in each of the null distributions. Finally, to assess whether decoding was above chance, we transformed the true decoding values to TFCE statistics and compared them to the 95^th^ percentile of the corrected null distribution.

### Behavioural data collection

In addition to our MEG experiment, we collected colour categorisation accuracies and reaction times on our stimuli from a new sample of 100 participants on Amazon’s Mechanical Turk. Participants were presented with the red and green shapes and the grey-scale objects, each presented individually for 100ms, randomly intermingled. On the instructions screen, participants were told that they would see images that can be categorised as red or green. They were informed that some images would be shown in greyscale, but that these objects were typically associated with red or green. Their task was to categorise the images into these two categories as fast and as accurately as possible by responding with either “m” or “z” using a keyboard. This allowed us to first confirm that the objects we had selected were indeed typically associated with red or green, and second, to test whether there was a reaction time difference between real and implied colour categorisation. Response-key mappings were randomly determined for each participant. Participants each completed 6 practice trials on objects that were not used in the experiment before the actual data collection began. Each participant was presented with each of the objects once. We calculated the mean accuracy and reaction times for the real and implied colour condition.

## Results

For our real colour decoding analysis, we trained the classifier to distinguish red from green shapes and then tested whether it could distinguish red from green shapes in an independent set of data. The classifier was able to predict the colour above chance in a cluster stretching from 65 to 315 ms, reflecting a signal modulated by colour (Figure 2, orange).

**Figure 2:**
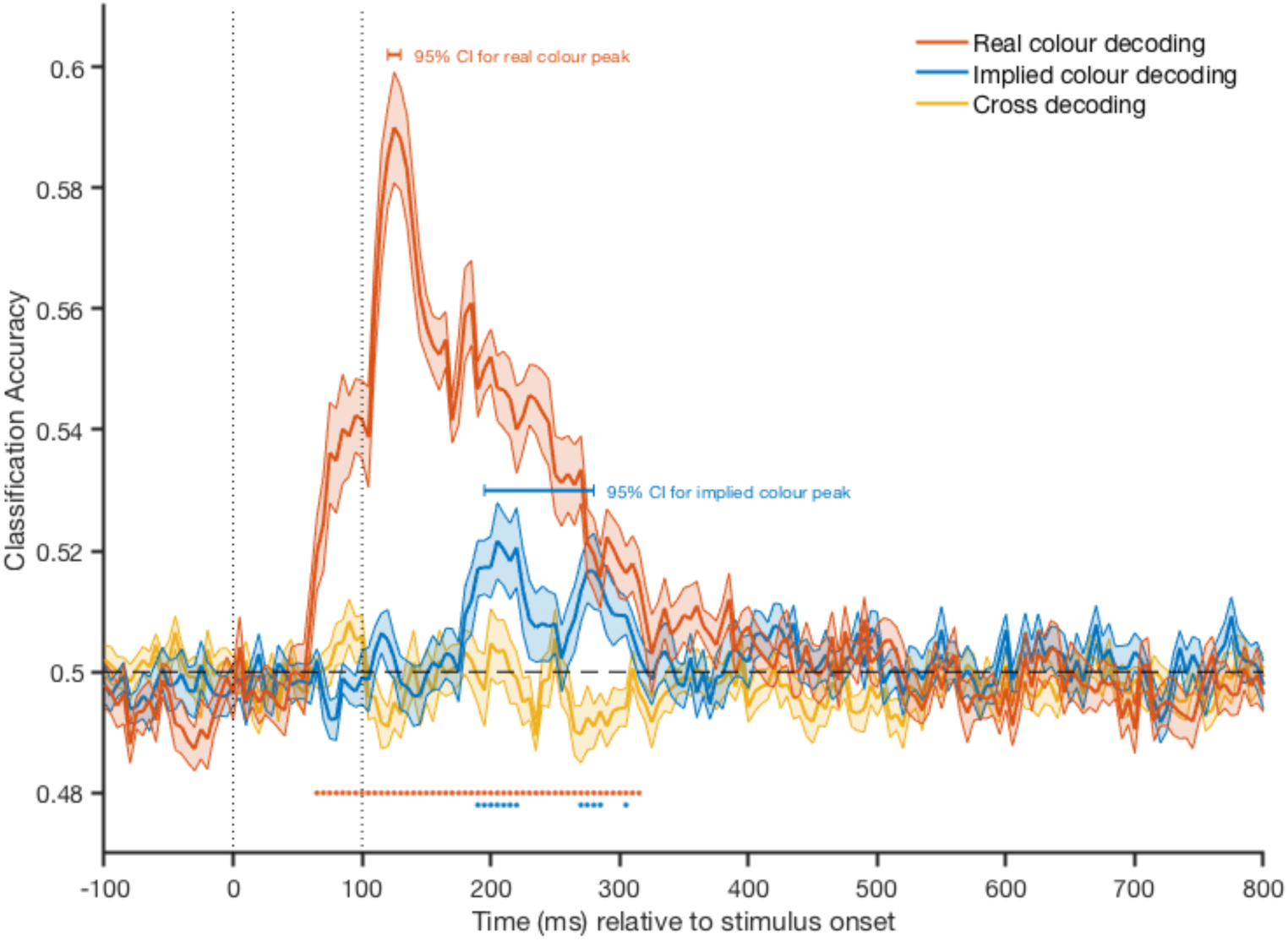
Classification accuracies for real colour (orange), implied colour (blue), and direct cross-colour (yellow) decoding over time. Vertical, dotted lines show stimulus on- and offset. Dashed line indicates chance level (50%). Shading indicates error bars around the mean (standard deviation of decoding accuracies across participants divided by the square root of the number of participants). Coloured dots depict significant timepoints corrected for multiple comparisons. The 95% confidence intervals for peak decoding latencies are plotted above the classification accuracies.

To examine whether we can decode implied object colour, the classifier was trained on a subset of the object trials and then tested on an independent set. The testing set included only exemplars (e.g., all strawberry and kiwifruit trials) that the classifier did not train on. Our data show that the classifier can distinguish between the objects belonging to the red and green category significantly above chance in a cluster stretching from 190 to 215 ms and from 270 to 290 ms (Figure 2, blue). While this suggests that there is categorical difference between objects associated with red and green, the results of this particular analysis could be driven by an overall difference in object characteristics other than colour (e.g., if red objects tend to have more round edges than green objects), and we therefore do not interpret this further.

Our key analysis to test whether there is a representational overlap of real and object colour processing depends on cross-decoding: training a classifier on real colour stimuli and testing on grey-scale objects that have implied colours. We trained the classifier to distinguish between the red and green shapes and tested its performance on the grey-scale objects to see whether direct cross-generalisation between real and implied object colour is possible. In this analysis, the classifier is trained on identical shapes that only vary in terms of colour. Hence, this is the most conservative way of testing whether there is a representational overlap between real and implied colours. The cross-colour decoding was not significant at any point in the timeseries (Figure 2, yellow). Accessing implied colour, however, presumably requires first accessing the general concept of the object. Therefore, real and implied colours may have a similar representation but colour information could be accessed later when activated via objects in comparison to when colour is perceived. We therefore tested whether this is the case using a cross-decoding time-generalisation analysis. We trained a classifier to distinguish between red and green shapes at every timepoint and then tested whether it could cross-generalise to the grey-scale objects at any timepoint. The results of key cross-generalisation analyses are summarised in Figure 3, showing a cluster of significant cross-generalisation with a time-shift.

**Figure 3:**
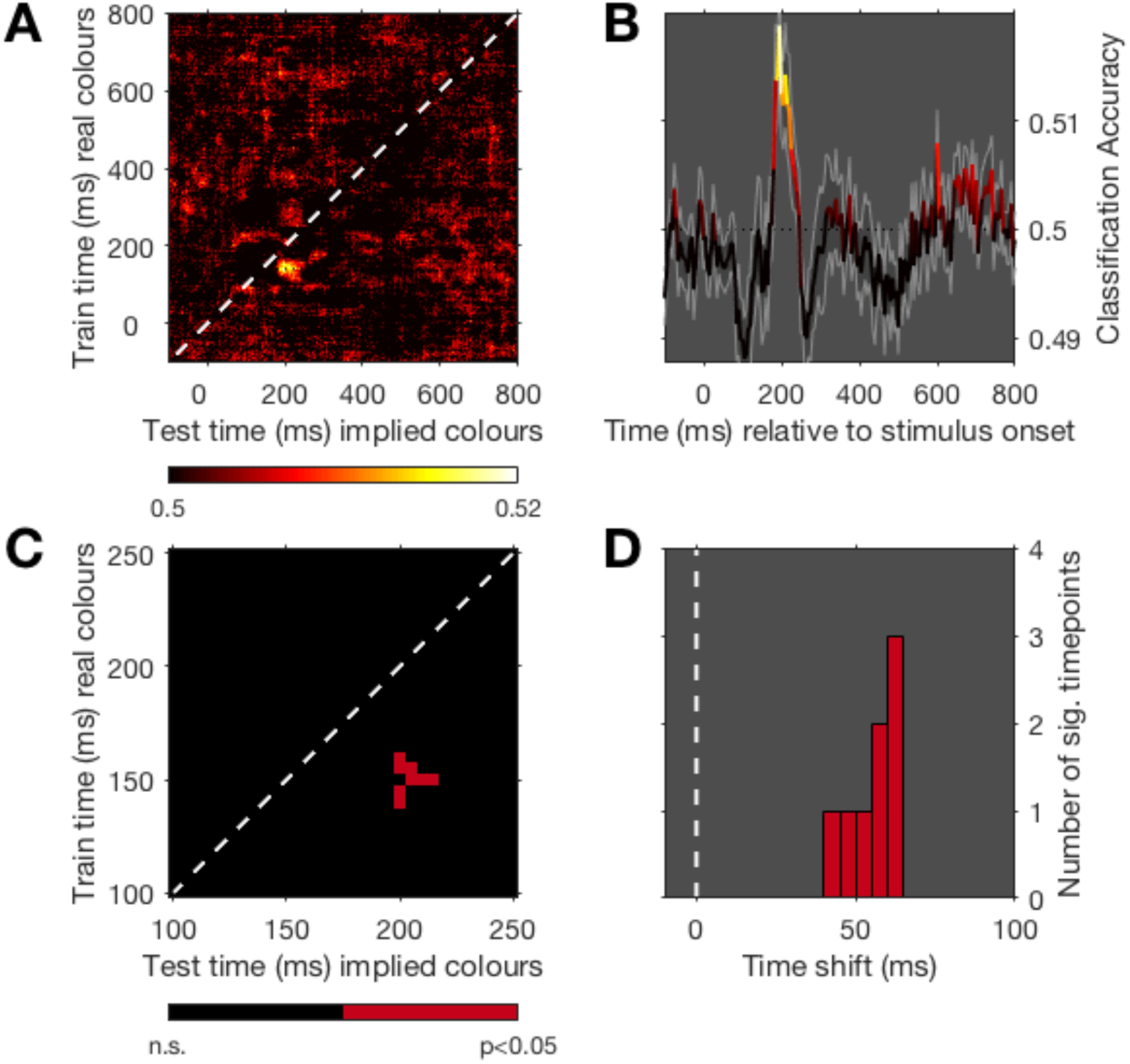
Results of the time-generalisation analysis. In this analysis, the classifier was trained on the real colour trials and tested on the implied colour trials. Panel A shows the classification accuracy at every possible train-test-time combination. Lighter colours indicate higher classification accuracy. Panel B shows the timecourse of the classification accuracy when the classifier relies on the training data at 140ms (peak decoding at 200ms). Shading indicates error bar around the mean (standard deviation of decoding accuracies across participants divided by the square root of the number of participants). Panel C shows all training-testing-timepoint combinations where classification is significantly above chance (based on permutation, corrected for multiple comparisons). Note that the axes in C are different to show the significant timepoints. Panel D shows the time shift of all significant timepoints from the diagonal. The delay of colour representations activated via implied colour activation in comparison to real colour perception is ∼55ms.

The time-generalisation analysis revealed similar activation patterns between real and implied colours when the classifier is trained on real colour at an earlier timepoint and tested on implied colour at a later one (Figure 3A and 3B). These generalisation accuracies were statistically above chance, even with our conservative correction for multiple comparisons (Figure 3C). Inspecting the training timepoint with maximum decoding (140 ms) indicates that there is above-chance decoding at later testing timepoints with peak decoding at 200 ms after stimulus onset (Figure 3B). The results show that we can cross-decode between real and implied colours when we *train* the classifier on *real* colours at timepoints between 140 to 160 ms and *test* it on *implied* colours at a cluster from 200 to 215 ms (Figure 3C). Combining the off-diagonal shift of the significant timepoints shows a median delay of 55 ms for implied colour testing times compared to real colour training times (Figure 3D). Importantly, these results are unlikely to be driven by anything else than colour as the classifier is trained on real colour trials in which the only different stimulus characteristic was colour and tested on implied colour trials which were achromatic. As a check, we also performed the reverse analysis (i.e., training the classifier on implied colour trials and testing it on real colour trials) which showed the same results, mirrored across the diagonal. The results highlight that there are similarities between real colour and implied object colour patterns but this pattern is instantiated later for implied object colours than for real colours. Note that above-chance cross-decoding does not mean we can interpret that the processes involved in real and implied colour processing are identical. However, the results show that there are sufficient similarities for the classifier to cross-generalise from brain activation patterns evoked by perceiving red and green to brain activation patterns evoked by viewing grey-scale images of objects that are associated with red and green^1^.

These results predict that it takes more time to access implied colour than real colour, presumably because one first has to access the concept of the object. We decided post-hoc to test this prediction behaviourally. 100 mTurk participants were presented with the red and green shapes and the grey-scale objects, each presented individually for 100ms, and were asked to indicate as quickly and accurately as possible whether the stimulus was (typically) red or green. Four participants were excluded from the analysis as their accuracy scores were more than 2 standard deviations below the group mean. For the remaining 96 participants, we excluded all the incorrect responses and compared the correct reaction times to the real and implied colour trials. Responding correctly to a real colour shape was on average ∼136ms (SD = 85ms) faster than responding correctly to an implied colour of an object (t(95) = 15.9, p<0.05, 95% CI [121.08, 155.64]). Real colour responses were also more accurate (M = 91.5%, SD = 0.08) than implied colour responses (M=80.4%, SD=0.13). The accuracy scores for the real and implied colour condition were significantly different (t(95)=8.07, p<0.05, 95% CI [8.3 13.8]). Using mTurk introduces variance to the experimental setup, including monitor settings for colour, computer and internet speeds, all of which will increase the noise in the data; we do not, therefore, interpret the specific difference in timing. Despite the variability, there is a clear difference between the time taken for categorising colour in the two conditions. These results are consistent with real colour perception being faster and easier than recalling implied colours, in line with the prediction from our decoding results.

To test the relationship between the neural data and behavioural data further, we also ran an exploratory analysis correlating the neural data of our sample with the behavioural data of the independent set of mTurk participants. We correlated the stimulus-wise behavioural categorisation data with the stimulus-wide MEG decoding accuracies for the implied colour decoding analysis and examined how this correlation unfolds over time (Figure 4). The results show that the neural data can be linked to the behavioural data from ∼200ms after stimulus onset which suggests that the information we decode can be used to generate behaviour (cf. de-Wit, Alexander, Ekroll, & Wagemans, 2016; Grootswagers, Cichy, & Carlson, 2018; Williams, Dang, & Kanwisher, 2007).

**Figure 4:**
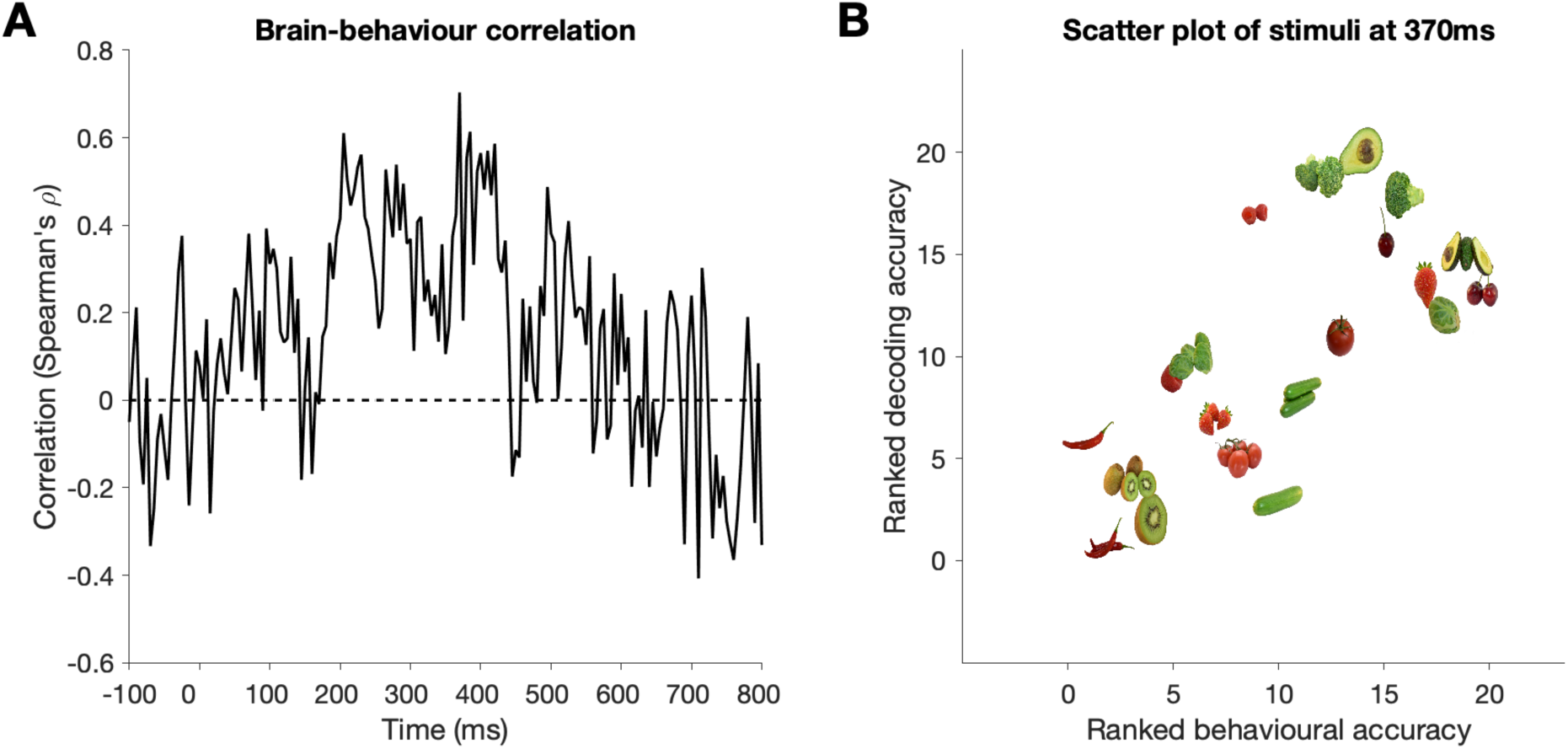
Panel A shows the correlation between the stimulus-wise behavioural accuracies in the independent colour categorisation task and the stimulus-wise MEG decoding accuracies over time. Panel B shows a scatterplot for the ranked behavioural accuracies and the ranked decoding accuracies for each stimulus at the peak timepoint of Panel A (370ms).

## Discussion

In this study, we compared the temporal activation patterns of colour perception and implied colour to examine the interaction between perceptual processing and access to object representations. We applied MVPA to time-series MEG data and show that both real and implied colour can be decoded, with some caveats around implied colour decoding due to potential visual stimulus differences. Our key results indicate that real and implied colour processing share a sufficient degree of similarity to allow for cross-generalisation with a temporal shift. The activity pattern distinguishing colours was instantiated ∼55ms later for implied colours than for real colour, highlighting that there are similarities between colour representations accessed via ‘real’ colour and via implied colour, but that there is a temporal asynchrony between these processes.

We interpret our cross-decoding results as evidence that the representation of implied colour involves some of the same mechanisms as those involved in colour perception. This is in line with previous studies showing that the same brain regions are active when retrieving object representations from memory and perceiving those object features (for reviews see A. Martin, 2007; Patterson et al., 2007) For example, Vandenbroucke et al. (2014) and Bannert and Bartels (2013) showed that early visual cortex is involved when real and implied colour are processed. Using fMRI, Vandenbroucke et al. (2014) trained a classifier on data recorded while participants viewed red and green shapes, and then tested the classifier on data recorded while participants viewed line-drawings of colour-diagnostic objects filled with ambiguous colours between red and green. Behavioural results suggested participants were slightly more likely to respond ‘red’ to the ambiguous colour presented on a line drawing of a typically red object than a line drawing of a typically green object. In their fMRI data, the classifier categorised the colour consistent with what the participant perceived. That means the classifier categorised colours to be red when shown on objects that are typically red, and green for objects that are typically green, at above chance levels. They interpret these data as evidence for an influence of implied object colours on the creation of a subjective experience of colour. Consistent with this study, Bannert and Bartels (2013) trained a classifier to distinguish fMRI data from trials where four different colours were presented. They showed that the classifier can cross-generalise to grey-scale colour-diagnostic objects. Both fMRI studies highlight that there are similar activation patterns across voxels in the visual cortex for real and implied colour processing. Our results provide further evidence that object-colour knowledge and colour perception instantiate similar patterns, this time in the temporal domain.

There are several possible explanations for a temporal difference between accessing colour representations via real colour and implied colour. One possibility is that the time difference reflects greater individual variability in the temporal activation profile of implied colours in comparison to real colours. Implied colours may be accessed at slightly different timepoints for different people and thus the cross-decoding accuracy that is above chance for each participant only overlaps at a later timepoint. There are also more interesting potential explanations. First, it could be due to actual differences in neural processes. Colour representations accessed via colour perception are immediately available whereas implied colour activation presumably only happens once the object is processed to some higher level. Thus, the delay could reflect differences between bottom-up and top-down access to colour representations. It might be, for example, that processing an object with a typical colour involves the activation of information about the object’s implied colour which is fed-back to earlier visual areas to compare incoming information with stored object-knowledge. In comparison, the shapes used in the real colour trials are not associated with a typical colour and thus do not evoke such as signal. This is a plausible interpretation of the temporal delay and corresponds with earlier findings of early visual areas being involved in implied colour activation (Bannert & Bartels, 2013; Vandenbroucke et al., 2014). Second, it is possible that the binding of colour and shape information happens later in the visual processing hierarchy compared to initial processing of separate features, and that the comparison of typical and perceived colour can therefore only happen later once shape-colour binding is complete. This view is consistent with results of a recent fMRI study, which showed that object-colour and object-shape activated from memory can be distinguished in areas associated with colour (V4) and shape (lateral occipital cortex, LOC) perception, respectively, but that the conjunction of colour *and* shape can be decoded only later along the visual hierarchy (anterior temopral lobe, ATL; Coutanche & Thompson-Schill, 2014). Similarly, Seymour et al. (2015) showed that colour *per se* can be decoded in early visual areas but object surface colour (bound to form) can only be decoded in areas further along the ventral visual stream. These findings also correspond to patient work (Patterson et al., 2007) and previous transcranial magnetic stimulation studies (Chiou, Sowman, Etchell, & Rich, 2014) which point towards the ATL as the hub for object-knowledge (for a review see Lambon Ralph, Jefferies, Patterson, & Rogers, 2017). Besides the ATL, other brain areas along the processing stream such as the medial temporal lobe (e.g., Rey et al., 2018), and the parahippocampal cortex (e.g., C. B. Martin, Douglas, Newsome, Man, & Barense, 2018) are also involved in retrieving long-term associations. Thus, it is possible that the temporal delay reflects the time it takes to activate these long-term colour associations. Finally, it could also be that the delay reflects the greater complexity of the grey-scale objects relative to the abstract shapes, hence binding the features may take slightly longer. From the data we cannot disentangle these interpretations. Our results clearly highlight, however, that there is a similar structure to the brain response to externally perceived and internally activated colour representations, and that time seems to be the key difference.

What is driving the successful decoding performance? For the real colour decoding, we used shapes that were identical across colour categories and used five different levels of stimulus luminance for each category that were perceptually matched. Therefore, the only distinguishing feature between the stimuli was colour. That means that for the real-colour decoding analysis *and* the cross-generalisation (i.e., training on shapes and testing on objects), we can rule out visual differences other than colour as a driving factor. Our results show that we can successfully decode real colour from ∼65ms onwards. The within-implied colour decoding results show that implied colour is decodable at ∼190ms after stimulus onset and then again a bit later at ∼270ms. This double-peak may occur because of variance between stimuli, such that accessing colour representations might be quicker for some images with stronger colour associations (for example) than others, or between participants in the speed with which they activate these representations. Alternatively, it may relate to differences in feedforward and feedback processes. For this within-implied colour classification analysis, visual differences could potentially contribute, as natural objects cannot be perfectly matched for the different conditions (i.e., red and green), unlike in our real colour condition. Previous studies have used line-drawings instead of photos of objects to reduce local low-level differences between stimuli (e.g., Vandenbroucke et al., 2014). Line-drawings can reduce some of these differences (e.g., local luminance differences due to texture) but also cannot completely rule out any contribution of low-level effects (e.g., shapes). In addition, there is a considerable trade-off between line-drawings in terms of similarity of the objects to real world objects which can slow down recognition and implied colour effects (Olkkonen, Hansen, & Gegenfurtner, 2008; Vurro, Ling, & Hurlbert, 2013). We therefore used isoluminant, grey-scale photos of objects and dealt with differences in low-level features (e.g., edges) by using an independent exemplar cross-validation approach. We trained the classifier to distinguish typically red and green objects using all objects except one typically-red and one typically-green object (each with two exemplars, which were both left out). The classifier was then tested on the left-out pair. We thereby considerably reduced the likelihood of the implied colour classification being driven by low-level features as the classifier never trained and tested on the same objects. While limiting the influence low-level features could have on the implied object colour decoding, it is still possible that the results in this particular analysis are driven by object features other than colour. To test this, we ran the same classification analysis on the output of a deep convolutional neural network which showed that it is unlikely that low-level visual differences account for all of the within-implied classification results (see supplementary material). Crucially, however, visual differences are not a concern for the key cross-decoding analysis. Here, we used identical red and green shapes in the training set, making low-level shape or texture features a highly unlikely source of contribution to classifier performance and colour hue being the primary predictor of category for the classifier (red vs green).

Our time-generalisation analysis shows that there are sufficient similarities in neural representation when perceiving real colour and activating implied colour for cross-generalisation. In addition, these results speak to the important aspect of temporal differences between colour evoked by external stimulation and internal activation. Activating conceptual knowledge of objects from memory is thought to involve a widespread network of brain areas involved in perception and action of different object features (A. Martin, 2007; Patterson et al., 2007). To access the implied colour of an object requires that the conceptual representation of that object is activated first. Using time-generalisation methods (King & Dehaene, 2014), we show here that in comparison to real colour perception, which can be decoded rapidly, accessing object-colour knowledge takes ∼55ms longer. This is consistent with our behavioural data showing that real colour judgments are faster than implied colour judgments. The behavioural data do not, however, speak to the neural similarity between real and implied colour activation patterns, which are observed in the time-generalisation analyses. Our MEG results increase our existing knowledge of how real and implied colour are processed by showing that aspects of colour representations via external stimulation are also instantiated during internal activation, but with a delay. Applying MVPA to our MEG data allows us to capture the similarity of representations of real colour perception and implied colour activation, but also allow us to examine temporal differences, highlighting the value of this method for dissociating activation of memory of object features from perception of object features in the real world.

Our results highlight that the activation of implied colours can occur independent of a task that focuses on colour. Participants completed a target-detection task in which attending to colour was not a useful strategy. The targets were ambiguous in colour (e.g., a capsicum can be either red or green), and this avoided biasing participants towards deliberately thinking about the implied colour of the objects. Using a task that is irrelevant to the classifier performance allowed us to explore the involuntary activation of implied colours rather than the signals associated with perhaps actively imagining colours or retrieving colour names. Not attending to the feature that is relevant for the classifier probably reduced our decoding accuracy in general (e.g., Brouwer & Heeger, 2013; Jackson, Rich, Williams, & Woolgar, 2017), but clearly supports previous studies showing that there is an *involuntary* activation of object-colour independent of task demands (Bannert & Bartels, 2013; Vandenbroucke et al., 2014).

Overall, the decoding accuracies across our analyses are low but significantly above chance with conservative statistics. As outlined above, this is probably partially due to colour being irrelevant for the task. In addition, it is important to note that we did not use extensive pre-processing, meaning we ran our analyses on effectively raw data. We use our multivariate decoding analyses for *interpretation* (Hebart & Baker, 2017)– if decoding is above chance, this means there is a signal that allows a categorical distinction between the conditions. Minimal pre-processing (e.g., no trial averaging, filtering, channel-selection, trial-selection) ensures that there is no potential influence of plurality of methods or specific pre-processing choices; it also means that the data overall are noisier which can result in relatively low decoding accuracies. However, it is crucial to note that low decoding accuracies does not necessarily mean that the effects are weak, as decoding accuracies are not effect sizes (cf. Hebart & Baker, 2017). Here, we show with rigorous methodological controls and strict correction for multiple comparisons that there is significant cross-generalisation from real colour to implied colour.

Previous fMRI studies showed that early visual areas are involved in real colour perception and implied colour activation (Bannert & Bartels, 2013; Rich et al., 2006; Vandenbroucke et al., 2014), but other studies implicate anterior temporal regions in object colour knowledge. For example, a transcranial magnetic stimulation study showed that the behavioural effects of implied colour knowledge on object recognition are disrupted when stimulating the anterior temporal lobe (Chiou et al., 2014), complementing patient studies suggesting this area holds conceptual object information (e.g., Lambon Ralph & Patterson, 2008). This highlights that activating object attributes, including implied colour, goes beyond low-level visual areas. Our study adds time as a novel aspect to this discussion by comparing the temporal profiles of colour representations accessed via real colour perception and implied colour activation.

In conclusion, our data show that there is a common representation of real and implied colour but that this representation is accessed later when triggered by activating implied colour than by perceiving real colour. This is in line with previous studies suggesting that the same brain areas are involved in object-feature activation from memory *and* object-feature perception. Our results highlight that applying MVPA to time-series MEG data is a valuable approach to exploring the interaction between object-feature inputs and predictions or representations based on prior knowledge. This opens multiple avenues for future studies examining the dynamic interactions between perceptual processes and activation of prior conceptual knowledge.

## ACKNOWLEDGEMENTS

This research was supported by the Australian Research Council (ARC) Centre of Excellence in Cognition and its Disorders, International Macquarie University Research Training Program Scholarships to LT & TG, an ARC Future Fellowship (FT120100816) and ARC Discovery project (DP160101300) to TC and an ARC Discovery project (DP170101840) to ANR.

## Footnotes

1 Please note that these results are stable across different analysis parameters. For example, the effect remains when using a different classifier, a wider sliding time windows, and when averaging across trials in the training data, normalising the training data, and using principal component analysis.

## References

Bannert, M. M., & Bartels, A. (2013). Decoding the yellow of a gray banana. Current Biology, 23(22), 2268–2272.

Barsalou, L. W., Simmons, W. K., Barbey, A. K., & Wilson, C. D. (2003). Grounding conceptual knowledge in modality-specific systems. Trends in Cognitive Sciences, 7(2), 84–91.

Bartleson, C. J. (1960). Memory colors of familiar objects. JOSA, 50(1), 73–77.

Bramão, I., Faísca, L., Petersson, K. M., & Reis, A. (2010). The influence of surface color information and color knowledge information in object recognition. The American Journal of Psychology, 123(4), 437–446.

Brouwer, G. J., & Heeger, D. J. (2013). Categorical clustering of the neural representation of color. Journal of Neuroscience, 33(39), 15454–15465.

Carlson, T. A., Hogendoorn, H., Kanai, R., Mesik, J., & Turret, J. (2011). High temporal resolution decoding of object position and category. Journal of Vision, 11(10), 9–9.

Chao, L. L., & Martin, A. (1999). Cortical Regions Associated with Perceiving, Naming, and Knowing about Colors. Journal of Cognitive Neuroscience, 11(1), 25–35. https://doi.org/10.1162/089892999563229

Chiou, R., & Rich, A. N. (2014). The role of conceptual knowledge in understanding synaesthesia: Evaluating contemporary findings from a “hub-and-spokes” perspective. Frontiers in Psychology, 5. https://doi.org/10.3389/fpsyg.2014.00105

Chiou, R., Sowman, P. F., Etchell, A. C., & Rich, A. N. (2014). A conceptual lemon: Theta burst stimulation to the left anterior temporal lobe untangles object representation and its canonical color. Journal of Cognitive Neuroscience, 26(5), 1066–1074.

Collins, J. A., & Olson, I. R. (2014). Knowledge is power: How conceptual knowledge transforms visual cognition. Psychonomic Bulletin & Review, 21(4), 843–860.

Coutanche, M. N., & Thompson-Schill, S. L. (2014). Creating concepts from converging features in human cortex. Cerebral Cortex, 25(9), 2584–2593.

de-Wit, L., Alexander, D., Ekroll, V., & Wagemans, J. (2016). Is neuroimaging measuring information in the brain? Psychonomic Bulletin & Review, 23(5), 1415–1428. https://doi.org/10.3758/s13423-016-1002-0

Engel, A. K., Fries, P., & Singer, W. (2001). Dynamic predictions: oscillations and synchrony in top–down processing. Nature Reviews Neuroscience, 2(10), 704.

Firestone, C., & Scholl, B. J. (2016). Cognition does not affect perception: Evaluating the evidence for “top-down” effects. Behavioral and Brain Sciences, 39.

Goldstone, R. L., de Leeuw, J. R., & Landy, D. H. (2015). Fitting perception in and to cognition. Cognition, 135, 24–29.

Grootswagers, T., Cichy, R. M., & Carlson, T. A. (2018). Finding decodable information that can be read out in behaviour. NeuroImage.

Grootswagers, T., Wardle, S. G., & Carlson, T. A. (2017). Decoding Dynamic Brain Patterns from Evoked Responses: A Tutorial on Multivariate Pattern Analysis Applied to Time Series Neuroimaging Data. Journal of Cognitive Neuroscience, 29(4), 677–697. https://doi.org/10.1162/jocn_a_01068

Hansen, T., Olkkonen, M., Walter, S., & Gegenfurtner, K. R. (2006). Memory modulates color appearance. Nature Neuroscience, 9(11), 1367.

Hebart, M. N., & Baker, C. I. (2017). Deconstructing multivariate decoding for the study of brain function. Neuroimage.

Hering, E. (1920). Grundzüge der Lehre vom Lichtsinn. Springer.

Jackson, J., Rich, A. N., Williams, M. A., & Woolgar, A. (2017). Feature-selective attention in frontoparietal cortex: multivoxel codes adjust to prioritize task-relevant information. Journal of Cognitive Neuroscience, 29(2), 310–321.

Kaiser, P. K. (1991). Flicker as a function of wavelength and heterochromatic ficker photometry. Limits of Vision, 171–190.

King, J. R., & Dehaene, S. (2014). Characterizing the dynamics of mental representations: the temporal generalization method. Trends in Cognitive Sciences, 18(4), 203–210. https://doi.org/10.1016/j.tics.2014.01.002

Kleiner, M., Brainard, D., Pelli, D., Ingling, A., Murray, R., Broussard, C., & others. (2007). What’s new in Psychtoolbox-3. Perception, 36(14), 1.

Lambon Ralph, M. A., Jefferies, E., Patterson, K., & Rogers, T. T. (2017). The neural and computational bases of semantic cognition. Nature Reviews Neuroscience, 18(1), 42.

Lambon Ralph, M. A., & Patterson, K. (2008). Generalization and differentiation in semantic memory: insights from semantic dementia. Annals of the New York Academy of Sciences, 1124(1), 61–76.

Maris, E., & Oostenveld, R. (2007). Nonparametric statistical testing of EEG-and MEG-data. Journal of Neuroscience Methods, 164(1), 177–190.

Martin, A. (2007). The representation of object concepts in the brain. Annu. Rev. Psychol., 58, 25–45.

Martin, A., Haxby, J. V., Lalonde, F. M., Wiggs, C. L., & Ungerleider, L. G. (1995). Discrete cortical regions associated with knowledge of color and knowledge of action. Science, 270(5233), 102–105.

Martin, C. B., Douglas, D., Newsome, R. N., Man, L. L., & Barense, M. D. (2018). Integrative and distinctive coding of visual and conceptual object features in the ventral visual stream. Elife, 7, e31873.

Olkkonen, M., Hansen, T., & Gegenfurtner, K. R. (2008). Color appearance of familiar objects: Effects of object shape, texture, and illumination changes. Journal of Vision, 8(5), 13–13.

Oostenveld, R., Fries, P., Maris, E., & Schoffelen, J.-M. (2011). FieldTrip: open source software for advanced analysis of MEG, EEG, and invasive electrophysiological data. Computational Intelligence and Neuroscience, 2011, 1.

Oosterhof, N. N., Connolly, A. C., & Haxby, J. V. (2016). CoSMoMVPA: multi-modal multivariate pattern analysis of neuroimaging data in Matlab/GNU Octave. Frontiers in Neuroinformatics, 10. Retrieved from https://www.ncbi.nlm.nih.gov/pmc/articles/PMC4956688/

Patterson, K., Nestor, P. J., & Rogers, T. T. (2007). Where do you know what you know? The representation of semantic knowledge in the human brain. Nature Reviews Neuroscience, 8(12), 976.

Pelli, D. G. (1997). The VideoToolbox software for visual psychophysics: Transforming numbers into movies. Spatial Vision, 10(4), 437–442.

Rey, H. G., De Falco, E., Ison, M. J., Valentin, A., Alarcon, G., Selway, R., … Quiroga, R. Q. (2018). Encoding of long-term associations through neural unitization in the human medial temporal lobe. Nature Communications, 9(1), 4372.

Rich, A. N., Williams, M. A., Puce, A., Syngeniotis, A., Howard, M. A., McGlone, F., & Mattingley, J. B. (2006). Neural correlates of imagined and synaesthetic colours. Neuropsychologia, 44(14), 2918–2925.

Seymour, K. J., Williams, M. A., & Rich, A. N. (2015). The representation of color across the human visual cortex: distinguishing chromatic signals contributing to object form versus surface color. Cerebral Cortex, 26(5), 1997–2005.

Simmons, W. K., Ramjee, V., Beauchamp, M. S., McRae, K., Martin, A., & Barsalou, L. W. (2007). A common neural substrate for perceiving and knowing about color. Neuropsychologia, 45(12), 2802–2810.

Smith, S. M., & Nichols, T. E. (2009). Threshold-free cluster enhancement: addressing problems of smoothing, threshold dependence and localisation in cluster inference. Neuroimage, 44(1), 83–98.

van Driel, J., Olivers, C. N., & Fahrenfort, J. J. (2019). High-pass filtering artifacts in multivariate classification of neural time series data. BioRxiv, 530220.

Vandenbroucke, A. R., Fahrenfort, J. J., Meuwese, J. D. I., Scholte, H. S., & Lamme, V. A. F. (2014). Prior knowledge about objects determines neural color representation in human visual cortex. Cerebral Cortex, 26(4), 1401–1408.

Vurro, M., Ling, Y., & Hurlbert, A. C. (2013). Memory color of natural familiar objects: Effects of surface texture and 3-D shape. Journal of Vision, 13(7), 20–20.

Williams, M. A., Dang, S., & Kanwisher, N. G. (2007). Only some spatial patterns of fMRI response are read out in task performance. Nature Neuroscience, 10(6), 685.

Witzel, C. (2016). An easy way to show memory color effects. I-Perception, 7(5), 2041669516663751.

