## Supplementary for "Seeing versus Knowing: The Temporal Dynamics of Real and Implied Colour Processing in the Human Brain"

**Supplementary Materials**

Table 1: HSV values for the colours used in the real experiment. Five consistent levels of luminance were used for red while green was perceptually matched to the reds for each participant with a colour flicker task. Hue and saturation were kept consistent throughout.

**Red: Five consistent levels of luminance**

| | Red1 | | | Red2 | | | Red3 | | | Red4 | | | Red5 | | |
|---|---|---|---|---|---|---|---|---|---|---|---|---|---|---|---|
| | H | S | V | H | S | V | H | S | V | H | S | V | H | S | V |
| | 0.00 | 0.90 | 0.70 | 0.00 | 0.90 | 0.77 | 0.00 | 0.90 | 0.84 | 0.00 | 0.90 | 0.91 | 0.00 | 0.90 | 0.98 |

**Green: Perceptually matched per participant**

| | Green1 | | | Green2 | | | Green3 | | | Green4 | | | Green5 | | |
|---|---|---|---|---|---|---|---|---|---|---|---|---|---|---|---|
| Participant | H | S | V | H | S | V | H | S | V | H | S | V | H | S | V |
| 1 | 0.333 | 0.9 | 0.405 | 0.333 | 0.9 | 0.435 | 0.333 | 0.9 | 0.48 | 0.333 | 0.9 | 0.51 | 0.333 | 0.9 | 0.505 |
| 2 | 0.333 | 0.9 | 0.405 | 0.333 | 0.9 | 0.455 | 0.333 | 0.9 | 0.48 | 0.333 | 0.9 | 0.47 | 0.333 | 0.9 | 0.52 |
| 3 | 0.333 | 0.9 | 0.405 | 0.333 | 0.9 | 0.425 | 0.333 | 0.9 | 0.45 | 0.333 | 0.9 | 0.495 | 0.333 | 0.9 | 0.49 |
| 4 | 0.333 | 0.9 | 0.4 | 0.333 | 0.9 | 0.475 | 0.333 | 0.9 | 0.39 | 0.333 | 0.9 | 0.455 | 0.333 | 0.9 | 0.475 |
| 5 | 0.333 | 0.9 | 0.44 | 0.333 | 0.9 | 0.475 | 0.333 | 0.9 | 0.485 | 0.333 | 0.9 | 0.52 | 0.333 | 0.9 | 0.56 |
| 6 | 0.333 | 0.9 | 0.525 | 0.333 | 0.9 | 0.525 | 0.333 | 0.9 | 0.53 | 0.333 | 0.9 | 0.535 | 0.333 | 0.9 | 0.56 |
| 7 | 0.333 | 0.9 | 0.39 | 0.333 | 0.9 | 0.415 | 0.333 | 0.9 | 0.42 | 0.333 | 0.9 | 0.475 | 0.333 | 0.9 | 0.495 |
| 8 | 0.333 | 0.9 | 0.385 | 0.333 | 0.9 | 0.415 | 0.333 | 0.9 | 0.515 | 0.333 | 0.9 | 0.49 | 0.333 | 0.9 | 0.485 |
| 9 | 0.333 | 0.9 | 0.415 | 0.333 | 0.9 | 0.43 | 0.333 | 0.9 | 0.445 | 0.333 | 0.9 | 0.475 | 0.333 | 0.9 | 0.48 |
| 10 | 0.333 | 0.9 | 0.395 | 0.333 | 0.9 | 0.415 | 0.333 | 0.9 | 0.435 | 0.333 | 0.9 | 0.47 | 0.333 | 0.9 | 0.45 |
| 11 | 0.333 | 0.9 | 0.45 | 0.333 | 0.9 | 0.43 | 0.333 | 0.9 | 0.495 | 0.333 | 0.9 | 0.485 | 0.333 | 0.9 | 0.47 |
| 12 | 0.333 | 0.9 | 0.42 | 0.333 | 0.9 | 0.455 | 0.333 | 0.9 | 0.48 | 0.333 | 0.9 | 0.51 | 0.333 | 0.9 | 0.55 |
| 13 | 0.333 | 0.9 | 0.42 | 0.333 | 0.9 | 0.425 | 0.333 | 0.9 | 0.415 | 0.333 | 0.9 | 0.455 | 0.333 | 0.9 | 0.455 |
| 14 | 0.333 | 0.9 | 0.5 | 0.333 | 0.9 | 0.47 | 0.333 | 0.9 | 0.48 | 0.333 | 0.9 | 0.505 | 0.333 | 0.9 | 0.565 |

| | | | | | | | | | | | | | | | |
|---|---|---|---|---|---|---|---|---|---|---|---|---|---|---|---|
| 15 | 0.333 | 0.9 | 0.46 | 0.333 | 0.9 | 0.445 | 0.333 | 0.9 | 0.485 | 0.333 | 0.9 | 0.525 | 0.333 | 0.9 | 0.52 |
| 16 | 0.333 | 0.9 | 0.455 | 0.333 | 0.9 | 0.52 | 0.333 | 0.9 | 0.495 | 0.333 | 0.9 | 0.54 | 0.333 | 0.9 | 0.545 |
| 17 | 0.333 | 0.9 | 0.41 | 0.333 | 0.9 | 0.445 | 0.333 | 0.9 | 0.43 | 0.333 | 0.9 | 0.47 | 0.333 | 0.9 | 0.475 |
| 18 | 0.333 | 0.9 | 0.43 | 0.333 | 0.9 | 0.455 | 0.333 | 0.9 | 0.575 | 0.333 | 0.9 | 0.615 | 0.333 | 0.9 | 0.57 |
| **MEAN** | **0.33** | **0.90** | **0.43** | **0.33** | **0.90** | **0.45** | **0.33** | **0.90** | **0.47** | **0.33** | **0.90** | **0.50** | **0.33** | **0.90** | **0.51** |

**Deep neural net classification**

It is possible that the within-decoding implied colour analysis could be influenced by low-level differences such as shape differences. We reduced the influence of shape on decoding accuracy by using a leave-one-exemplar-out cross-validation, but the most important thing is that the key finding (cross-decoding) cannot be explained by these differences, as the classifier trains on the shape trials which only differ in terms of colour and tests on completely different achromatic stimuli.

In addition to being very cautious in our interpretation of the within-decoding implied colour results, we test whether differences in the images could drive our decoding analyses by repeating these analyses using the activation of units in the VGG-19 deep neural network. Here, we plot the decoding performance for each layer:

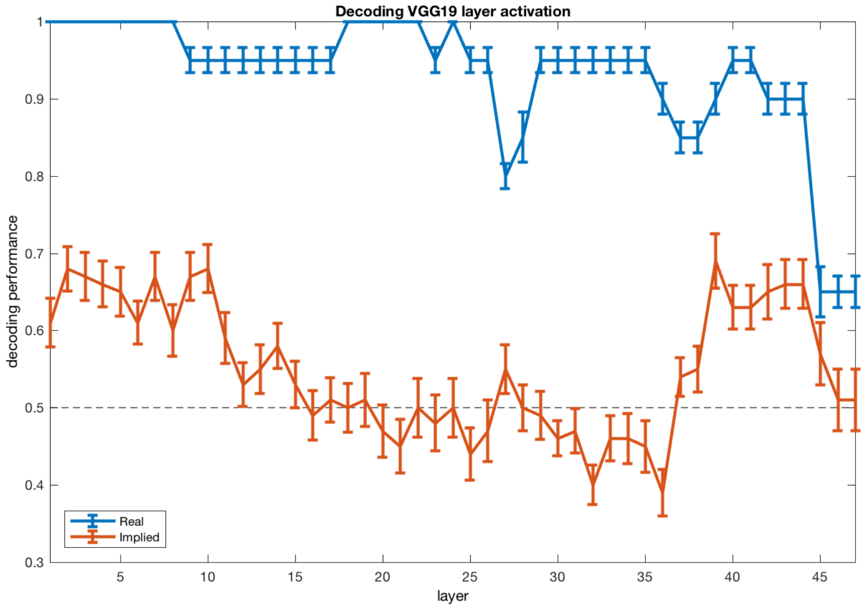


Supplementary Figure 1: Decoding VGG19 layer activation

Similar to our MEG decoding results, decoding performance across all layers was superior for real colours in comparison to implied colours. For the implied colour decoding analysis, there is a peak in the early layers (1-10) and then again in later layers (from 37 onwards).

The peak in the early layers suggests that part of the implied colour decoding from our MEG data could be driven by visual features. To examine this possibility further, we used the DNN output and plotted the correlation between the objects. We visualised the level of similarity using multidimensional scaling (MDS) across all layers (an example for Layer 4 is shown in Figure 2 below).

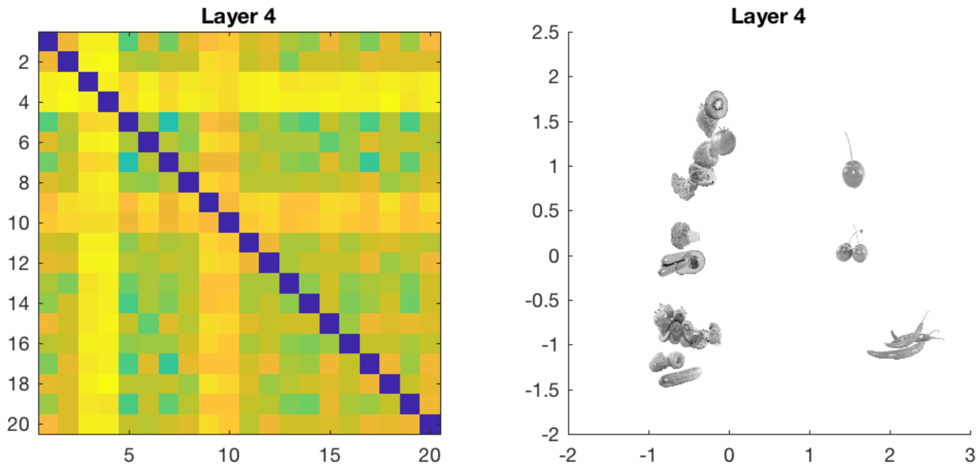


Supplementary Figure 2: Dissimilarity matrix and multidimensional scaling results for one layer (4) for the grey-scale objects.

If the MEG decoding performance is driven by the visual features that the DNN can distinguish, we should be able to see that the same objects that are driving the classifier performance in the DNN (i.e., cherries and chillies, Supplementary Figure 2) also drive the classifier performance in the MEG data. We therefore looked at the MEG decoding accuracies for each of the objects individually at the significant timepoints. Although cherries and chillies contribute to the MEG decoding accuracy, they are not the only contributors (Supplementary Figure 3). This suggests that the classifier did not use the same information as the DNN to categorise implied colours from the MEG data.

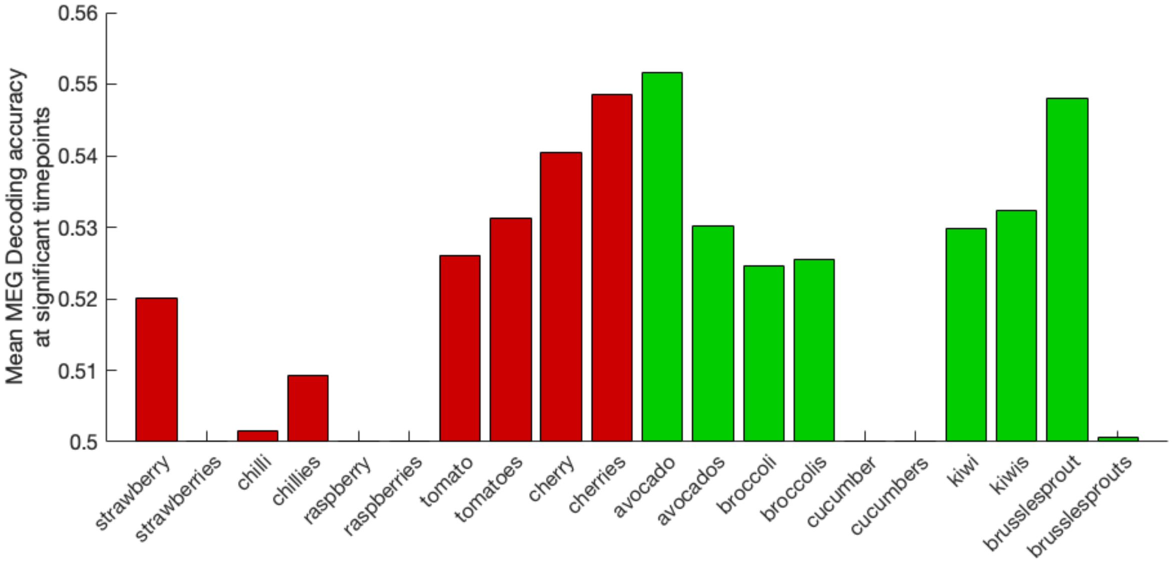


Supplementary Figure 3: Mean MEG decoding accuracy for each object averaged across the significant timepoints of the implied-colour decoding analysis.
